# Characterisation of β-tubulin isotypes in *Uncinaria stenocephala* and implications for benzimidazole resistance in hookworms

**DOI:** 10.1101/2025.07.12.664546

**Authors:** Thomas Stocker, Jan Šlapeta

**Affiliations:** Sydney School of Veterinary Science, Faculty of Science, The University of Sydney, New South Wales 2006, Australia; Sydney Infectious Diseases Institute, New South Wales 2006, Australia

**Keywords:** *tubb-1*, *tubb-2*, whole genome sequence, WGS, mutation, exon, Ancylostomatidate

## Abstract

*Uncinaria stenocephala* is a widespread hookworm of dogs across Europe, Canada, southern Australia, and other temperate regions, where it often outnumbers infections caused by *Ancylostoma caninum*. Although a putative β-tubulin isotype-1 mutation associated with resistance has been detected in *U. stenocephala*, clinical resistance to benzimidazoles has not yet been confirmed. Benzimidazole resistance is primarily linked to single nucleotide polymorphisms (SNPs) in the β-tubulin isotype-1 gene; however, the β-tubulin genes of *U. stenocephala* have not been fully characterised. We aimed to identify β-tubulin genes and confirm the coding sequences for key residues (Q134, F167, E198, and F200) in the β-tubulin isotype-1 gene of the *U. stenocephala* genome. Two *U. stenocephala* specimens were subjected to Illumina sequencing, and species identity was confirmed through morphological and molecular analysis using ITS rDNA and cox-1 markers. Genome assembly revealed the presence of β-tubulin isotype-1 (10 exons) and isotype-2 (9 exons), both homologous to β-tubulins from other hookworms (*A. caninum*, *A. ceylanicum*, *A. duodenale* and *Necator americanus*). The β-tubulin isotype-1 protein sequence of *U. stenocephala* contained two variable residues (S37Q and G441A) compared to other hookworm sequences. While the isotype-2 protein sequence was conserved among *Ancylostoma* species, *U. stenocephala* exhibited six distinct polymorphisms (E39D, T40S, N115S, L130I, A287S, T439G). The benzimidazole-susceptible residues (Q134, F167, E198, F200) were present in the β-tubulin isotype-1 protein sequence. Characterisation of the complete coding regions of β-tubulin isotypes 1 and 2 enables population-level screening for benzimidazole resistance-associated SNPs and provides a foundation for future epidemiological studies in *U. stenocephala*.

**Highlights:** β-tubulin isotypes-1 and -2 were fully characterised in *Uncinaria stenocephala*

Key benzimidazole-susceptible residues were confirmed in β-tubulin isotype-1

β-tubulin isotypes-1 and -2 enables SNP screening and resistance surveillance efforts

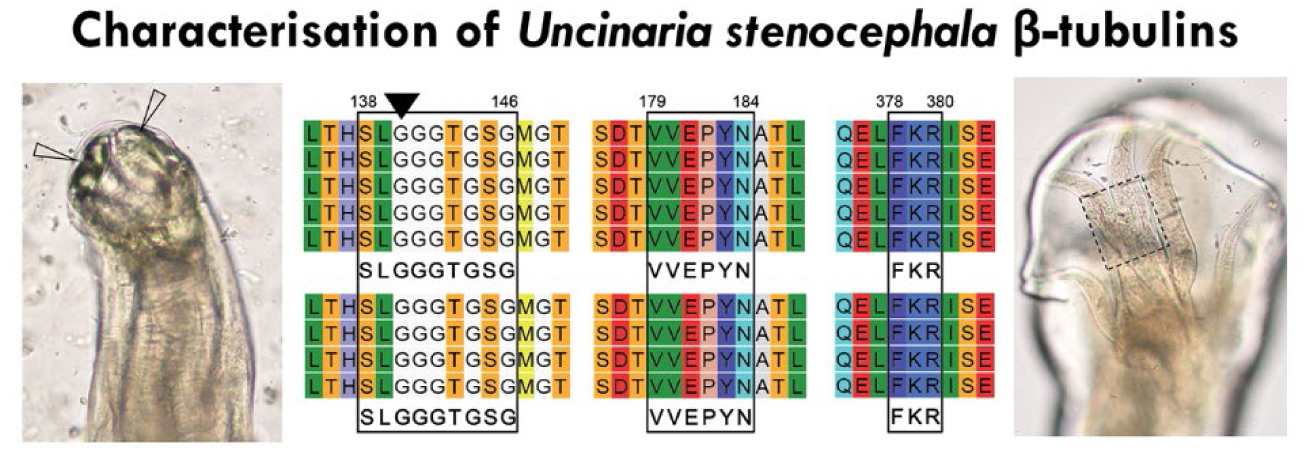

## 1. Introduction

Hookworm nematodes (Ancylostomatidae) cause anaemia in humans, cats, and dogs through gastrointestinal haemorrhages. Until recently, however, anthelmintic resistance had not been a major concern (Geary et al., 2025). Benzimidazoles have been used effectively for over 50 years in companion animals to treat hookworm infections. Their mode of action involves disrupting the assembly of the nematode cytoskeleton and impairing metabolism, phenotypically visible through disrupted development. Clinical resistance to benzimidazoles was first documented in the canine hookworm (*Ancylostoma caninum*) using both *in vivo* (faecal egg count reduction test, FECRT) and *in vitro* (egg hatch assay, EHA) methods (Jimenez Castro et al., 2019; Kitchen et al., 2019). While FECRT and EHA are invaluable, they are inherently low-throughput. High-throughput amplicon sequencing has emerged as a powerful alternative for screening known single nucleotide polymorphisms (SNPs) in tubulin genes, which are critical components of the nematode cytoskeleton. Amplicon metabarcoding and sequencing of the β-tubulin isotype-1 (*tubb-1*) gene have revealed widespread mutations at amino acid positions 134 and 167, which confer benzimidazole resistance in *A. caninum* populations from dogs in North America and Australia (Abdullah et al., 2025; Venkatesan et al., 2023).

While mutations in the *tubb-1* gene have been strongly associated with benzimidazole resistance in hookworms, the broader genetic context of β-tubulin genes in nematodes warrants further investigation. Strongyle nematodes, including hookworms, possess four genes encoding β-tubulins (*tubb-1*, *tubb-2*, *tubb-3* and *tubb-4*), in contrast to the model organism *Caenorhabditis elegans*, which also contains the *ben-1* gene and is amenable to genetic studies (Collins et al., 2024; Saunders et al., 2013). Although non-synonymous variants at codons 134, 167, 198 and 200 of the *tubb-1* gene have been associated with benzimidazole resistance, *tubb- 1* may not be the sole gene involved (Collins et al., 2024). The causal role of these codon changes was demonstrated through mutations in the *ben-1* gene of *C. elegans*, which confer resistance to benzimidazoles (Dilks et al., 2020; Dilks et al., 2021; Kitchen et al., 2019; Venkatesan et al., 2023). In *C. elegans*, *ben-1* confers the highest level of resistance, and the loss of a second β-tubulin gene does not alter this resistance level (Collins et al., 2024). In nematodes lacking *ben-1*, resistance is conferred by other β-tubulin genes, with *tubb-1* playing a major, though not exclusive, role. Therefore, it is important to define the full complement of β- tubulin genes, particularly *tubb-1* and *tubb-2*, which are ubiquitously expressed in nematode cells, and to screen for mutations that may either confer or contribute to benzimidazole resistance (Collins et al., 2024; Gandasegui et al., 2024; Saunders et al., 2013; Venkatesan et al., 2023). Currently, no additional β-tubulin isotypes (*tubb-3* and *tubb-4*) have been identified in hookworms.

Hookworm nematodes are among the common intestinal parasites of companion animals, particularly dogs and cats, alongside roundworms and whipworms (Lightner et al., 1978). In these hosts, the primary pathogenic hookworms are *A. caninum* in dogs and *Ancylostoma tubaeforme* in cats (Miller, 1968). However, in cooler temperate regions, *Uncinaria stenocephala*, traditionally considered less pathogenic, predominates. Unlike the blood-feeding *Ancylostoma* species that cause anaemia, *U. stenocephala* infections are associated with protein-losing enteropathy (Walker and Jacobs, 1985). Recognised as the ‘northern hookworm’, *U. stenocephala* is widely distributed across Europe, Canada, southern Australia, and New Zealand, where it often outnumbers *Ancylostoma* spp. infections (Beveridge, 2002; Gibbs, 1961; Xhaxhiu et al., 2011).

Given the recent evidence of benzimidazole resistant-associated mutation in *U. stenocephala*, a deeper molecular characterisation of its β-tubulin genes is needed. Building on molecular tools developed for *A. caninum*, Stocker et al. (2023) demonstrated that a *tubb-1*- targeting assay could also amplify *U. stenocephala tubb-1*, making it suitable for screening potential genetic markers of benzimidazole resistance in this species. A study from Australia reported the presence of the canonical F167Y benzimidazole resistance mutation in *U. stenocephala tubb-1* (Abdullah et al., 2025). Although clinical treatment failure with benzimidazoles in *U. stenocephala* has not yet been documented, proactive molecular surveillance is essential. To anticipate emerging resistance, comprehensive characterisation of β-tubulin genes, particularly full-length *tubb-1*, across hookworm species is critical for enabling comparative analyses and early detection of resistance-associated mutations.

The aim of this study was to characterise the *tubb-1* and *tubb-2* genes of *U. stenocephala* and to identify key amino acid residues potentially associated with benzimidazole resistance. To achieve this, we isolated adult *U. stenocephala* specimens and performed whole-genome sequencing. Using a comparative approach with available hookworm genomic data from *A. caninum*, *Ancylostoma ceylanicum*, *Ancylostoma duodenale*, and *Necator americanus*, we identified and characterised the coding sequences of *tubb-1* and *tubb-2*. To facilitate high-throughput screening of benzimidazole-susceptible residues in TUBB-1 (Q134, F167, E198, and F200), we reviewed existing primer sequences and in silico assessed their suitability for *U. stenocephala* and other hookworm species.

## 2. Materials and Methods

### 2.1. Adult Uncinaria stenocephala material

To obtain *Uncinaria stenocephala*, we decided to consider the invasive red fox (*Vulpes vulpes*) a non-native species with significant ecological impact in Australia and is regularly culled (Stobo-Wilson et al., 2021). Red foxes are commonly infected by *U. stenocephala* in Australia (Ryan, 1976). Single red fox intestinal tract was donated by a registered shooter in September 2024 (Mardi; 33.2990° S, 151.4084° E); the fox was culled under an ongoing fox control program in Central Coast region of New South Wales, Australia. The gastrointestinal tract was transported on ice to the University of Sydney, stored for 10 days at -80°C prior to dissection. During dissection the gastrointestinal track was thawed, incised longitudinally and content scraped through the researcher’s thumb and index finger. The scrapings and small intestinal contents washed with lukewarm tap water into a 150 µm nylon mesh sieve by means of water jet. The washed content was examined in a white metal tray with the aid of stereomicroscope. Nematodes were manually picked against the white tray, washed in phosphate buffered saline (pH=8.0) and stored at -80°C. Hookworms, such as *U. stenocephala*, were identified in accordance with the description of Ransom (1924).

### 2.2. Genomic DNA extraction and molecular identification

Genomic DNA (gDNA) was isolated using the Monarch® Spin gDNA Extraction Kit (New England Biolabs, Australia). The gDNA was stored at -20°C. The isolated DNA was subject to two separate PCR assays. The PCR amplifying ITS rDNA used NC16 [S0863] (5’- AGT TCA ATC GCA ATG GCT T -3’) and NC2 [S0864] (5’- TTA GTT TCT TTT CCT CCG CT -3’) (Chilton et al., 2003) The partial *cox*1 was amplified with JB3 [S0361] (5’- TTT TTT GGG CAT CCT GAG GTT TA -3’) and JB4.5 [S0362] (5’- TAA AGA AAG AAC ATA ATG AAA ATG -3’) (Bowles et al., 1992). Amplicons of expected size (∼800 bp, ITS rDNA; ∼450 bp, *cox*1) were submitted for Sanger sequencing (Macrogen, Korea). Sequences were assembled and compared to available sequences in CLC Main Workbench 25.0.1 (CLC bio, Qiagen, Australia). As a reference of *U. stenocephala* ITS rDNA sequence we used (Nadler et al., 2000) and *cox*1 we used AJ407939 (Hu et al., 2002).

### 2.3. Whole genome sequencing and assembly

The gDNA obtained from two single *U. stenocephala* was aliquoted (30 μL) into a 1.5 mL Eppendorf tube and submitted to Novogene (HK) Co., Ltd for indexing, library preparation and whole genome sequencing to generate approximately 30 Gb of sequencing data per sample. Sequencing was performed using the Illumina NovaSeq 6000 sequencing platform using 150 bp paired-end sequencing chemistry. Raw FastQ files (JS6871 and JS6872) generated from the same sample were processed in Galaxy Australia (https://usegalaxy.org.au/) (Galaxy Community, 2024). Raw paired-end sequencing reads were examined for quality using *FastQC* (version 0.12.1) (Andrews, 2010) and trimmed using *Trimmomatic* (version 0.36.6) (Bolger et al., 2014) (parameters: “ILLUMINACLIP:TruSeq3-PE.fa:2:30:10 SLIDINGWINDOW:4:20”). Read were mapped using *bwa-mem2* (version 2.2.1) (Li and Durbin, 2009) using simple Illumina mode onto red fox genome (VulVul3) and all mapper reads were removed. The FastQ files were assembled using *SPAdes* (version 4.2.0) (Prjibelski et al., 2020) (parameters: “--cov-cutoff 40.0 -k 77”). Contigs were classified using *Kraken* (version 1.1.1) (database: minikraken) (Wood and Salzberg, 2014) and only unclassified contigs were retained. Assembly statistics were generated using *Fasta Statistics* (Seemann and Gladman, 2012).

### 2.4. Identification of β-tubulin genes in the genome assemblies

The *tubb-1* and *tubb-2* genes and their corresponding proteins were retrieved from hookworm genomes at ‘WormBase ParaSite’ (Howe et al., 2016; Howe et al., 2017). The genes were identified in *A. caninum, A. ceylanicum, A. duodenale* and *Ne. americanus* with and addition of an outgroup (*Ni. braziliensis*) (International Helminth Genomes, 2019; Schwarz et al., 2015). The reference sequences for *tubb-1* (ANCCAN_08153, Acey_s0035.v2.g13228, ANCDUO_05511, Necator_chrII.pre1.g8352 and NBR_0000799801) and *tubb-2* (ANCCAN_23405, Acey_s0432.v2.g1106, Necator_chrII.pre1.g8352 and Nippo_chrII.g5566) were aligned and genes structure identified. There was no *tubb-2* for *A. duodenale* localised due to partial coverage of the genome.

Local BLAST databases from the assembled contigs of *U. stenocephala* were created and queried with the reference *tubb-1* and *tubb-2* genes sequences. Candidate contigs were aligned to the reference alignments and exon-intron boundaries identified. Alignments and local BLAST searches were performed in CLC Main Workbench v 25.0.1 (CLC bio, Qiagen, Australia).

Sequence reads were mapped to the putative *tubb-1* and *tubb-2* using *bwa- mem2* (version 2.2.1) (Li and Durbin, 2009) using simple Illumina mode. *Samtools mpileup* (Version 1.15.1) (Danecek et al., 2021) was used to input variants into *VarScan* (Version 2.4.2) (Koboldt et al., 2012) (parameters: --min-coverage 20 --min-reads2 2 --min-avg-qual 15 --min-var-freq 0.2 --min-freq-for-hom 0.75 --p-value 0.05). The resulting BAM and VCF files were visualised using *IGV* (version 2.18.1). VCF file was processed to select only SNP within the exons of *tubb-1* and *tubb-2.* The variant detection was performed in Galaxy Australia (https://usegalaxy.org.au/) (Galaxy Community, 2024).

To evaluate the binding of the primers targeting *A. caninum tubb-1* (ACB1_167_F: 5’-GGY GCA GGA AAC AAC TG -3’, ACB1_167_R: 5’-CTT TGG TGA GGG GAC AAC A -3’, ACB1_200_F: 5’-GTR GTG GAG CCA TAC AAT GC -3’ and ACB1_200_R: 5’-GGC ATG AAG AAG TGA AGA CGT -3’) we first mapped them onto the reference *A. caninum tubb-1* and then in an multiple sequence alignment that included the *U. stenocephala tubb-1* we evaluated the number of mismatches across the primer binding region. The primers were originally development for amplicon metabarcoding to consider the frequency of the SNPs that code for the nonsynomynous mutations at 134, 167, 198 and 200 amino acid residue of *A. caninum tubb-1* responsible for benzimidazole resistance (Jimenez Castro et al., 2019; Venkatesan et al., 2023).

### 2.5. Data availability

The *U. stenocephala* sequences were submitted to GenBank under the following accession numbers: PV848420 and PV848421 (ITS rDNA), PV848422 and PV848423 (*cox*1), PV133906 (*tubb-1*) and PV862047 (*tubb-2*). Raw FastQ sequence data were deposited at SRA NCBI BioProject: PRJNA1281746 (https://www.ncbi.nlm.nih.gov/bioproject/PRJNA1281746). The additional data are available at LabArchives (https://dx.doi.org/10.25833/ffcf-2175).

## 3. Results

### 3.1. Morphological and molecular confirmation of *Uncinaria stenocephala*

Four hookworms (5-7 mm) were extracted from the small intestinal contents of a red fox (*V. vulpes*, TSF_1). The hookworms (2 males, 2 females) were consistent with the morphology of *U. stenocephala* (**Figure 1**; Ransom (1924))

**Figure 1:**
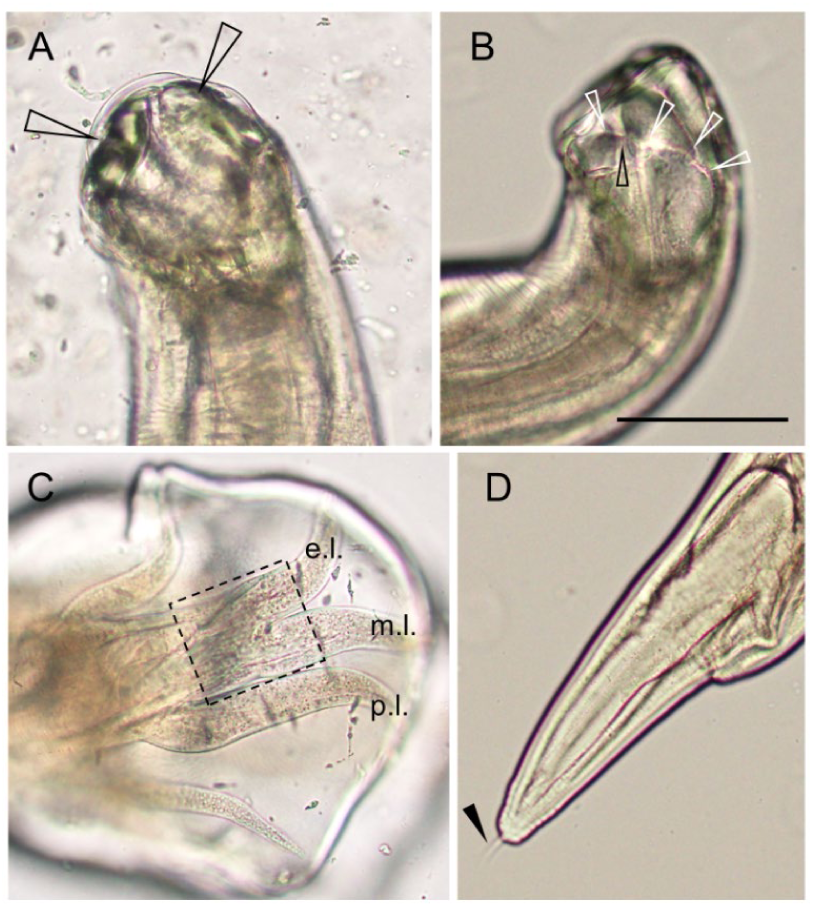
Morphology of sampled *Uncinaria stenocephala*. (A) Dorso-ventral view of the anterior end. Bicuspid cutting plates are visualised with an arrow for each cutting plate. (B) The lateral view of the anterior end. The internal cephalic papilla is labelled with a black arrow. The white arrows signify the boundary line between the thick ventral portion and the thinner dorsal portion of the lateral wall of the oral capsule. This boundary crosses through the internal cephalic papilla which is consistent with *U. stenocephala*. (C) Male copulatory bursa with the medio-lateral (m.l.) ray is roughly equal in width to the externo-lateral ray (e.l.) and the postero-lateral ray (p.l.). (D) Female posterior end with a caudal bristle (arrowhead). Scale bar for A-D , 100 μm.

Two *U. stenocephala* (TSF_1.1, TSF_1.2) were molecularly characterised at ITS rDNA and *cox*1 gene. Sequenced ITS rDNA (818 nt) were identical to each other and to the reference sequence of *U. stenocephala* (AF194145). Comparison of the new *U. stenocephala* ITS rDNA with *U. stenocephala* ITS2 rDNA data from amplicon metabarcoding data from dog faeces (Abdullah et al., 2025) revealed 95.8% to 100% match across 260 nt. The sequenced partial *cox*1 (393 nt) sequence from the two *U. stenocephala* differed by a single synonymous mutation (99.8% identity), and they were 99.2-99.5% identical to the reference *U. stenocephala* sequence (AJ407939).

### 3.2 Sequencing and assembly of *Uncinaria stenocephala* genome

Sequencing of TSF_1.1 (JS6871) and TSF_1.2 (JS6872) generated 117,718,741 and 102,660,825 paired-end reads (150 bp), respectively, with average GC contents of 46% and 45%. This yielded a total of 35.2 Gb for TSF_1.1 and 30.6 Gb for TSF_1.2. After quality trimming, 117,358,176 (99.7%) and 102,353,896 (99.7%) high-quality paired-end reads were retained. There were 4% and 23% sequence reads that mapped to the red fox (VulVul3) genome that were removed from TSF_1.1 and TSF_1.2, respectively. The final datasets comprised 31.0 Gb for TSF_1.1 and 21.6 Gb for TSF_1.2, both with a GC content of 45-46%. Each sample was assembled independently. The TSF_1.1 assembly had N50 of 2,203 bp and L50 of 32,460 contigs. We then removed potential bacterial or viral contaminant contigs (TSF_1.1: 0.2% of the contigs) to produce final assembly contigs (TSF_1.1: N50 of 2,090 bp and L50 of 32,766 contigs) (**Table 1**). The TSF_1.2 assembly had N50 of 8,683 bp and L50 of 2,153 contigs, following decontamination it had N50 of 8,431 bp and L50 of 2,065 contigs (**Table 1**).

**Table 1.** Assembly of the whole genome sequencing of *Uncinaria stenocephala*.

|  | TSF_1.1 | TSF_1.1* | TSF_1.2 | TSF_1.2* |
| --- | --- | --- | --- | --- |
| L50 | 32,460 | 32,766 | 2,152 | 2,065 |
| N50 | 2,203 | 2,090 | 8,683 | 8,431 |
| Number of bp | 341,439,219 | 333,472,904 | 62,113,770 | 57,967,112 |
| Number of sequences | 1,039,638 | 1,037,453 | 50,108 | 49,593 |
| GC content (%) | 45.5 | 45.6 | 44.6 | 44.7 |

\* putative bacterial and viral contaminants removed via Kraken

The TSF_1.1 assembly yielded a contig (NODE_5499; length 7,645 nt, coverage 19.8) that contained the complete *tubb-1* gene using blast searches with other *tubb-1* from hookworms as a query. The *tubb-2* was represented by three contigs, one contig (NODE_15465; length 4,291 nt, coverage 30.3) included all but the first exon and was truncated at the first intron. The first exon with partial first intron was found on two contigs, where the first exon was 100% identical between the contigs (NODE_82476; length 558 nt, coverage 29.6; NODE_38650; length 1,780 nt, coverage 24.1). The *tubb-2* complete coding sequence was fused from the contigs, the nucleotide sequence is truncated in the first exon. There were no contigs that would contain the complete putative *tubb-1* and *tubb-2* in the TSF_1.1 assembly.

### 3.3 Characterisation of tubb-1 and tubb-2 of Uncinaria stenocephala

The putative *tubb-1* and *tubb-2* of *Uncinaria stenocephala* genes were used to reference map the available sequence reads. A total of 1,774 and 1,256 reads were successfully mapped to the *tubb-1*, resulting in an average coverage of 59.7 and 41.6 bases per position across the covered regions from TSF_1.1 and TSF_1.2, respectively. Similarly, a total of 2,335 and 1,610 reads were successfully mapped to the *tubb-2*, resulting in an average coverage of 68.7 and 47.2 bases per position across the covered regions from TSF_1.1 and TSF_1.2, respectively.

The *U. stenocephala tubb-1* gene was 4,011 nt long including 10 exons homologous to other *btub-1* form hookworms (**Figure 1A**). The largest intron between the exon 4 and 5 was 1,688 nt, shorter compared to the intron from *A. caninum tubb-1* (46 nt: ANCCAN_08153 and 1217 nt: DQ459314). The *U. stenocephala btub-2* gene was 4649 nt long and had 9 exons homologous to those of other hookworms (**Figure 1B**). VarScan has detected 9 independent heterozygous nucleotide position in TSF_1.1 and TSF_1.2 within *tubb-1* of *U. stenocephala*. Within *tubb-2* of *U. stenocephala* there were 48 and 32 heterozygous positions in TSF_1.1 and TSF_1.2, respectively.

The coding protein sequence of *U. stenocephala* TUBB-1 had two amino acid changes when compared to other hookworms (S37Q and G441A), the remaining animo acid sequence was identical with other hookworms. The TUBB-2 protein of *U. stenocephala* had five amino acid changes compared to the TUBB-2 from *Ancylostoma* species: E39D, T40S, N115S, L130I and A287S. In addition, there is a single change shared with *N. americanus* TUBB-2 (T439G). Both TUBB-1 and TUBB-2 proteins included tubulin and β-tubulins protein motifs: VPRAVLVDLEP, SLGGGTGSG, VVEPYN and the β-tubulin specific motif FKR (**Figure 1C**).

We then compared the primers designed by (Jimenez Castro et al., 2019) for the amplification of the canonical SNPs in *tubb-1* of *A. caninum* with that of *U. stenocephala* (**Figure 2A**). The ACB1_167_F primer sequence was a perfect match, while ACB1_167_R had single, G>A, mismatch in the middle of the primer sequence compared to the *U. stenocephala tubb-1*. Both ACB1_200_F and ACB1_200_R had single mismatch each compared to the *U. stenocephala tubb-1*. The mismatch in the ACB1_200_R is at the 3’ terminal nucleotide position, T>C.

**Figure 2:**
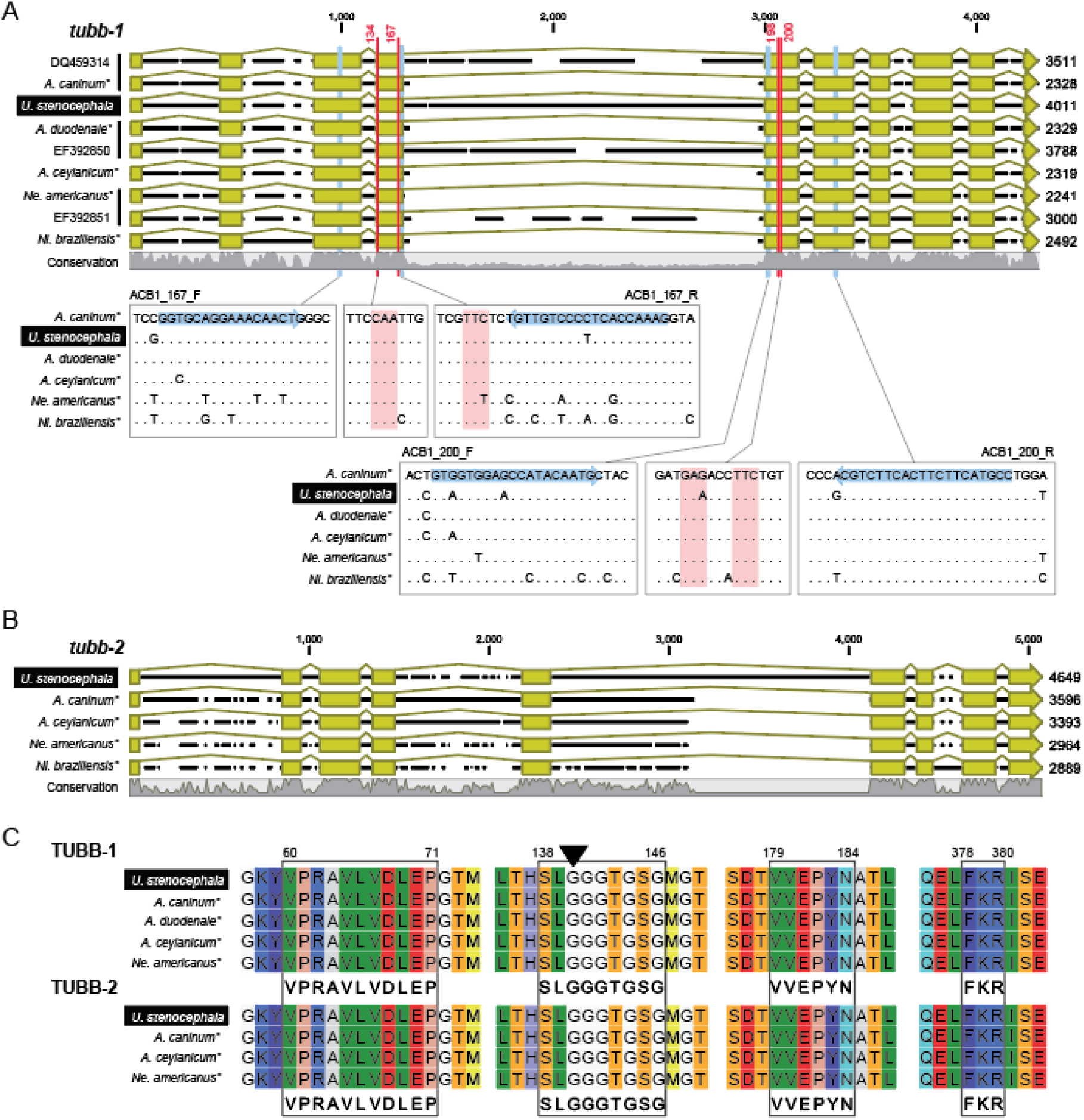
Comparison of *Uncinaria stenocephala* structure of *tubb-1* and *tubb-2* genes with other hookworms. (A) Gene sequence showing exon-intron boundaries of *tubb-1* of hookworms. Regions where primers targeting *A. caninum tubb-1* (ACB1_167_F: 5’-GG**<u>Y</u>** GCA GGA AAC AAC TG -3’, ACB1_167_R: 5’-CTT TGG TGA GGG GAC AAC A -3’, ACB1_200_F: 5’-GT**<u>R</u>** GTG GAG CCA TAC AAT GC -3’ and ACB1_200_R: 5’-GGC ATG AAG AAG TGA AGA CGT -3’) and benzimidazole-susceptible residues in *tubb-1* (Q134, F167, E198, and F200) are enlarged to show nucleotide sequences (identical residues to the top sequence as dots). (B) Gene sequence showing exon-intron boundaries of *tubb-2* of hookworms. Names with (*) indicate that sequence data were obtained from WormBase Parasite, *tubb-1* (ANCCAN_08153, Acey_s0035.v2.g13228, ANCDUO_05511, Necator_chrII.pre1.g8352 and NBR_0000799801) and *tubb-2* (ANCCAN_23405, Acey_s0432.v2.g1106, Necator_chrII.pre1.g8352 and Nippo_chrII.g5566), otherwise sequences are labelled by GenBank accession number. (C) Amino acid alignment regions of TUBB-1 and TUBB-2 showing the tubulin motifs VPRA(V/I)(F/L)(V/L)DLEP, S(F/L/A)AGGTGSG and VV(E/N)PYN motifs. TUBB-1 second motive devaiates from the standard motive (A140G, arrowhed). The β-tubulin conserved protein motive F(K/R)R motif in preset across all sequences.

## 4. Discussion

We successfully characterised the coding sequences of two representatives of the β-tubulin gene family in *U. stenocephala*. We show, that both the *tubb-1* and *tubb-2* genes are highly conserved among hookworms and retain an identical exon–intron structure. Access to the complete *U. stenocephala tubb-1* sequence enabled us to confirm the presence of key residues (Q134, F167, E198, and F200) that are associated with benzimidazole susceptibility in hookworms (Venkatesan et al., 2023). The corresponding TUBB-1 protein sequence was found to be homologous to the partial *U. stenocephala* sequences previously identified in amplicon metabarcoding studies (Abdullah et al., 2025; Stocker et al., 2023; Stocker et al., 2024). Anthelmintic resistance in companion animals is an emerging concern, and in the canine hookworm *A. caninum*, it has become an established reality (Geary et al., 2025; von Samson-Himmelstjerna et al., 2021). This study represents the first successful sequencing of the full *tubb-1* gene in the northern hookworm *U. stenocephala*. Characterisation of both *tubb-1* and *tubb-2* now enables population-level screening for resistance-associated SNPs in *U. stenocephala*, similar to the foundational work for *A. caninum* by (Schwenkenbecher et al., 2007).

The European red fox is a widespread invasive species in Australia (Stobo-Wilson et al., 2021). The prevalence of *U. stenocephala* in red foxes was reported to range from 18.2% to 30.6% (Coman, 1973; Dybing et al., 2013; Ryan, 1976). Using ITS rDNA and partial *cox*1 sequences, we demonstrated that *U. stenocephala* isolates from red foxes are genetically indistinguishable from those found in domestic dogs (Abdullah et al., 2025; Górski et al., 2023). Given that wild red foxes are not exposed to anthelmintic treatments, their high infection rates could represent a substantial refugium for benzimidazole-susceptible hookworms. However, the detection of the *tubb-1* F167Y SNP, associated with benzimidazole resistance, in *U. stenocephala* from an urban dog in Australia by Abdullah et al. (2025) suggests that the epidemiology of this parasite in dogs and foxes may be more complex and distinct than currently assumed.

Although there is currently no conclusive clinical evidence of benzimidazole treatment failure in *U. stenocephala*, the detection of the F167Y SNP in the *tubb-1* gene by Abdullah et al. (2025) provides early warning of potential resistance. This finding suggests that the conditions for resistance are present, and it may only be a matter of time before the frequency of this SNP increases to levels associated with clinical drug failure. In the absence of phenotypic validation of the F167Y SNP as a resistance marker in *U. stenocephala*, we must remain vigilant and anticipate the spread of benzimidazole resistance, as has occurred in *A. caninum* populations in Australia and North America (Abdullah et al., 2025; Geary et al., 2025; Leutenegger et al., 2024; Leutenegger et al., 2023; Venkatesan et al., 2023).

DNA polymerase fidelity is highly sensitive to mismatches at the 3’ end of PCR primers, particularly when the 3’ terminal nucleotide is incorrectly base-paired, as this can hinder the initiation of extension by the enzyme (Kwok et al., 1990). In the case of the ACB1_200_R primer, a T>C mismatch at the 3’ end poses a potential risk for PCR failure due to impaired primer extension. However, previous studies have shown that this mismatch did not interfere with successful amplification of the *U. stenocephala tubb-1* amplicon using the ACB1_200_F/ACB1_200_R primer pair (Abdullah et al., 2025; Stocker et al., 2023; Stocker et al., 2024; Venkatesan et al., 2023). With our characterisation of the *U. stenocephala tubb-1* gene and comparative analyses with other hookworm species, there is now an opportunity to refine these primers, either through degeneracy or redesign, to better accommodate sequence variability. The availability of genome sequencing data and expanded hookworm reference datasets further enhances the potential for improved diagnostic accuracy and data reliability in high- to medium-throughput molecular applications (Ilík et al., 2024).

We concluded that the identified protein sequnces belong to *U. stenocephala* β-tubulin, specifically TUBB-1 and TUBB-2. Burns (1991) established three protein motifs common to all tubulins, the VPRA(V/I)(F/L)(V/L)DLEP, S(F/L/A)AGGTGSG and VV(E/N)PYN motifs. These motifs were detected in all TUBB-1 and TUBB-2 from hookworms, with a sinlge exception. There was a deviation at the third residue of the second motif (A140G) aparently shared across all hookworms. All β-tubulin posses the motive F(K/R)R motif that was conserved across all hookworms. Whereas α- and γ-tubulin have KFD or RKR motif instead, respectively (Burns, 1991). Burns (1991) considered the hypervariable regions of the tubulin genes to infer functional residues. The divergent TUBB-1 residues and four of the six divergent TUBB-2 residues (E39D, T40S, A287S, T349G) of *U. stenocephala* were located in hypervariable regions and so unlikely to impact the function of the β-tubulins. The benzimidazole-susceptible residues Q134, F167, E198, and F200 were present in *U. stenocephala* TUBB-1 and were homologues to those in other hookworm sequences.

## 5. Conclusion

Benzimidazole- and multidrug-resistant *A. caninum* are present in domestic dog populations across North America and Australia (Abdullah et al., 2025; Kitchen et al., 2019; Venkatesan et al., 2023). In contrast, although *U. stenocephala* is the predominant hookworm species in canines from cooler regions such as continental Europe, southern Australia, and likely other under-sampled areas, data on its distribution and resistance profile remain scarce. By characterising the complete coding regions of *tubb-1* and *tubb-2*, we enable population-level screening for benzimidazole resistance-associated SNPs and lay the groundwork for epidemiological studies in the ‘northern hookworm’ *U. stenocephala*. Genetic screening revealed widespread SNPs in *A. caninum tubb-1* linked to benzimidazole resistance, indicating that these alleles circulated within populations long before clinical drug failures were recognised by veterinary parasitologists (Geary et al., 2025). Geary et al. (2025)recently proposed a set of action points to address resistance in *A. caninum*; we argue that these recommendations are equally relevant and urgently needed for both *A. caninum* and *U. stenocephala*.

## Acknowledgements

We acknowledge the Coastal Wildlife Management, Australia, that for donating the red fox material. T. Stocker would like to acknowledge the support and encouragement from Megan and Heidi. The computational part was supported by Galaxy Australia, a service provided by Australian BioCommons and its partners; the service receives NCRIS funding through Bioplatforms Australia, as well as The University of Melbourne and Queensland Government RICF funding.

## Author contributions: CRediT

**Thomas Stocker:** Conceptualization, Formal analysis, Investigation, Writing – original draft. **Jan Šlapeta**: Conceptualization, Funding acquisition, Project administration, Validation, Writing – review and editing.

